# XlinkCyNET: a Cytoscape application for visualization of protein interaction networks based on cross-linking mass-spectrometry identifications

**DOI:** 10.1101/2020.12.20.423654

**Authors:** Diogo Borges Lima, Ying Zhu, Fan Liu

**Affiliations:** Department of Chemical Biology, Leibniz - Forschungsinstitut fu◻r Molekulare Pharmakologie (FMP), Berlin, Germany

**Keywords:** protein-protein interaction, structural biology, cross-linking mass spectrometry, Cytoscape, bioinformatics

## Abstract

Software tools that allow visualization and analysis of protein interaction networks are essential for studies in systems biology. One of the most popular network visualization tools in biology is Cytoscape, which offers a large selection of plugins for interpretation of protein interaction data. Chemical cross-linking coupled to mass spectrometry (XL-MS) is an increasingly important source for such interaction data, but there are currently no Cytoscape tools to analyze XL-MS results. In light of the suitability of Cytoscape platform but also to expand its toolbox, here we introduce **XlinkCyNET**, an open-source Cytoscape Java plugin for exploring large-scale XL-MS-based protein interaction networks. XlinkCyNET offers rapid and easy visualization of intra and intermolecular cross-links and the locations of protein domains in a rectangular bar style, allowing subdomain-level interrogation of the interaction network. XlinkCyNET is freely available from the Cytoscape app store: http://apps.cytoscape.org/apps/xlinkcynet and at https://www.theliulab.com/software/xlinkcynet.

## INTRODUCTION

Proteins are essential building blocks of nearly all cellular processes. They interact with each other by forming stable protein complexes or transiently via signaling nodes, ensuring the diverse and precise function of the cell^1^. So far, several software tools have been developed to display large interaction networks using node-link diagrams, such as Biogrid^2^ and Cytoscape^3^. In particular, Cytoscape, as a popular software for network visualization and analysis, offers a great selection of plugins capable of interpretating network data from different angles. For instance, Cerebral^4^ provides options in Cytoscape to layout the network based on subcellular locations of proteins, thereby facilitating the analysis of the subcellular arrangements of signaling pathways. Another example is stringApp^5^, which integrates a major protein network database, STRING^6^, into Cytoscape, allowing visualization and analysis of networks retrieved by STRING using Cytoscape features (*e.g.*, enrichment analysis).

An increasingly important method for the systematic investigation of protein interaction networks is cross-linking mass spectrometry (XL-MS). In XL-MS, native protein contacts are captured using a cross-linker, which typically contains two functional groups that are reactive to specific amino acids. Cross-linked proteins are then subjected to enzymatic digestion and the resulting cross-linked peptides are analyzed by tandem mass spectrometry (MS/MS). These experiments yield a large number (often in the range of hundreds to tens of thousands) of residue-to-residue connections that can be used to generate protein interaction maps and to probe protein binding interfaces^7^. XL-MS data can be graphically visualized as network graphs using web-server applications (*e.g.*, XVis^8^ and xiNET^9^) or standalone software tools (*e.g.*, SIM-XL^10,11^ and Cross-ID^12^). Similar to Cytoscape, these applications present protein interactions using node-link diagrams. Despite successful applications in small and medium-sized networks, their effectiveness is often compromised in proteome-wide XL-MS studies due to the crowdedness of the display. Furthermore, while these XL-MS-specific tools allow users to import customized protein annotations (*e.g.*, proteins domains and post-translational modifications), they cannot compete with the versatility of the Cytoscape app store, as their features are often tailored to specific types of analysis, limiting the integration with other existing analysis tools.

To overcome these limitations and in light of the suitability of Cytoscape platform, we introduce XlinkCyNET, a Cytoscape app that supports not only the visualization of residue-to-residue connections of proteins provided by XL-MS data, but also protein domain annotations. As XlinkCyNET is fully integrated into Cytoscape, it connects the XL-MS-based network to all available features in Cytoscape, which allows users to take full advantage of the software to increase the quality of the analysis.

## METHODS

### XlinkCyNET development

XlinkCyNET is programmed in Java™ SE Development Kit (JDK) 11.0.8 utilizing the Cytoscape 3.8 Application Programming Interface (API). It is compatible with operating systems including Windows, Linux and MacOS. The app is an open-source project at GitHub and freely available to any interested party for further development. XlinkCyNET fulfils two aims: (1) display residue-to-residue cross-links in Cytoscape using a rectangular bar style and (2) allow annotation of protein domains in the network.

### Workflow

An overview of the XlinkCyNET workflow is depicted in **Figure 1**. XlinkCyNET allows two input files: (1) A mandatory network file containing essential information of cross-linked residues and proteins, and (2) an optional file that supplements domain annotation of proteins in the network. The loaded network is graphically displayed in the main *Network View Window* and the dataset is presented in the *Table Panel* of Cytoscape.

**Figure 1.**
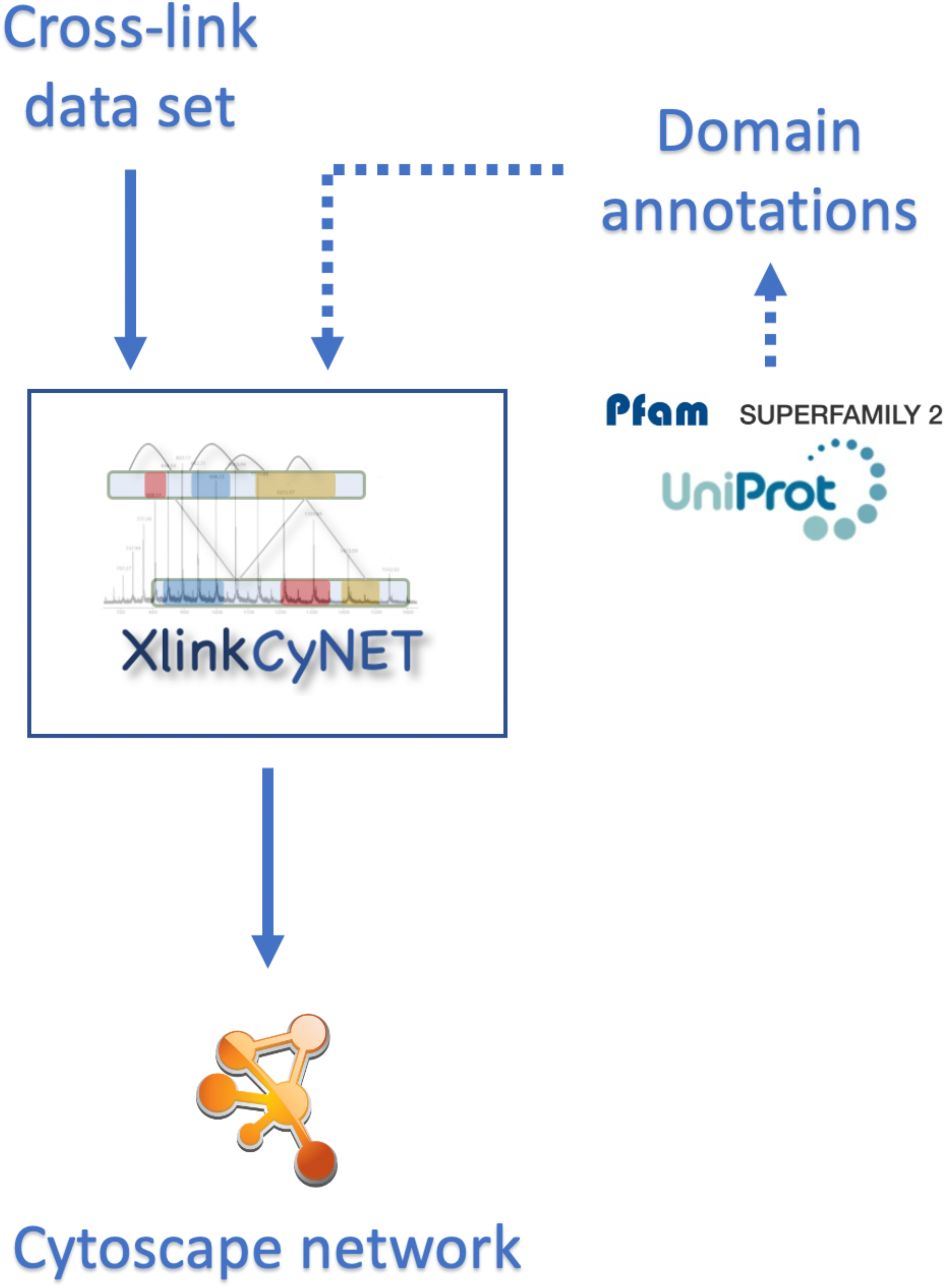
Overview of XlinkCyNET workflow. Cross-link dataset is a required input (solid arrow) file to guide Cytoscape in applying the reside-to-residue layout in the network. Domain annotation is an optional file (dash arrow) and can be imported to annotate domains of each protein.

### Input and output

XlinkCyNET supports input from all file types accepted by Cytoscape, such as, Simple Interaction Format (SIF), PSI-MI, SBML, XML-based BioPAX, GML, JSON, MS Excel™ Worksheets and comma- or tab-separated files (*i.e.*, *CSV* or *tab* files). To present cross-linked residues, XlinkCyNET requires the input file containing the following information: node identifier (*e.g.*, gene name, protein name), protein length and cross-links involved in each protein-protein interaction pair. In the use cases presented here, *CSV* files were used as input.

For protein domain annotation, XlinkCyNET accepts *CSV* files with the following pattern: names of different domains and ranges of amino acids in separated columns. XlinkCyNET also recognizes *tab* files that are directly downloaded from UniProtKB^**13**^. In this case, the *tab* file must contain *Gene names* (in *Names & Taxonomy category*) and *Topological domain* (in *Subcellular location category*) columns. Examples of different input files are shown in **Table S1** and **S2** respectively. For the output, the displayed network can be exported in all formats supported by Cytoscape (*e.g.*, JPG, SVG, PS and PDF). Protein domains modified directly in Cytoscape can be exported as *CSV* files via XlinkCyNET export function.

### Features and settings

The main features of XlinkCyNET are under *Apps* **→** *XlinkCyNET* (**Figure 2a**). Here all protein domains can be imported and exported via *Protein domains* tab. Protein domains can be loaded via two options: (1) upload an input file via *Protein domains* **→** *Load* **→** *File* **→** *Import file*, or (2) directly download the information from the SuperFamily^**14,15**^ (also known as Supfam) Distributed Annotation System server^[1]^ or the Pfam^**16**^ web-server^[2]^ via *Protein domains* **→** *Load* **→** *Search for domains* (**Figure 2b**). User-defined colors can be set for specific domains under *Protein domains* **→** *Set domain color* option.

**Figure 2.**
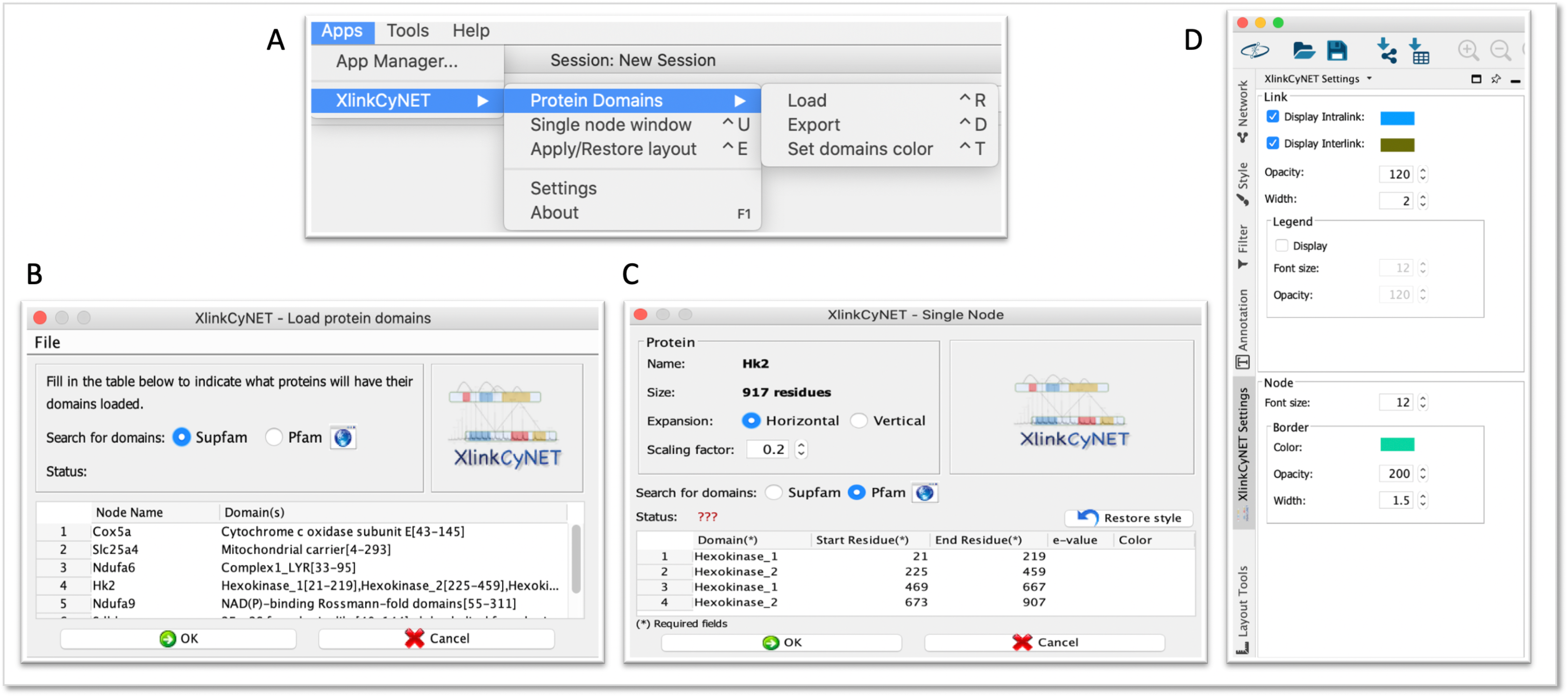
Overview of XlinkCyNET settings. (A) XlinkCyNET main menu. (B) *Load protein domains* window (*XlinkCyNET* → *Protein Domains* → *Load*) which supports uploading customized domain annotation file and downloading information from Supfam and Pfam webservers. (C) *Single Node* window (*XlinkCyNET* → *Single node window*) which shows all adjustable parameters for each protein. (D) *Settings* panel (*XlinkCyNET* → *Settings)* which provides different display options for cross-links, proteins and labels.

*Single node window* opens a pop-up window containing additional parameters of a single selected protein (**Figure 2c**). Here some parameters for individual proteins can be modified. For instance, *Scaling factor* allows the adjustment of the protein length presented as rectangular bars; *Expansion* determines the direction of the expansion as horizontal or vertical. Both parameters are stored in *Node Table*, shown in *scaling_factor* and *is_horizontal_expansion* columns, respectively. Domain information of each protein can be manually inserted in *Domain table* or downloaded from Supfam and Pfam webservers. The original Cytoscape layout can be restored by clicking on *Restore style* button.

*Apply/Restore layout* button applies the rectangular bar layout to all selected proteins in the network. A second click on *Apply/Restore layout* will restore the original Cytoscape style to all selected proteins. This action can be performed to both single and multiple proteins.

XlinkCyNET main settings are under *Settings*, where various display options (*e.g.*, color, width, font size and opacity) can be selected for cross-links, proteins (shown as rectangular bars) and labels (**Figure 2d**). As built-in functions of Cytoscape, cross-link sites are displayed when the mouse moves over the connections. Modifications of visualization parameters of a protein, such as fonts, colors and domain annotations, are updated in the main *Network View Window* when the protein is moved once.

### Program design

XL-MS experiment produces undirected cross-links, therefore, they are represented by unidirectional ed es in node-link diagrams. For inter-protein cross-links (*i.e.*, cross-links between two proteins), the algorithm computes the distance between the beginning of the protein and the location of the cross-linked residue based on the information provided by the input dataset. This method allows XlinkCyNET to draw and position a straight line (represented by a Cytoscape edge) on the rectangle bar in the exact position of the cross-linked site. For intra-protein cross-links (*i.e.*, cross-links within one protein), the algorithm considers the distance between the two cross-linked sites to compute the arc of the circumference. The arc is also represented by a Cytoscape edge, however, a *bend* property is applied to curve the edge. XlinkCyNET uses multi-threading to increase the p speed.

## RESULTS AND DISCUSSION

### XlinkCyNET for displaying large interaction networks: analysis of XL-MS data of the mitochondria interactome

We selected a previously published mitochondrial XL-MS dataset^17^ to demonstrate the functionality of XlinkCyNET. This dataset contains 359 proteins and 3,322 unique reside-to-residue contacts. Here we use XlinkCyNET version 1.1 and Cytoscape version 3.8.2 for illustration. We first formatted the XL-MS data provided in Supplementary Table 1 of the original publication using an *in-house* R-script and generated a cross-link-containing *CSV* file as the input (**Table S2**). We then imported the data using Cytoscape *from Import Network From File* function. By default, the network is displayed with only protein-protein interactions, whereas the residue-to-residue connections provided by XL-MS are hidden (Error! Reference source not found.**a**). Using Cytoscape *New Network From Selected Nodes, All Edges* function, we highlighted proteins in the mitochondrial oxidative phosphorylation complexes (OXPHOS) and extracted all associated nodes and edges to a subnetwork (**Figure 3b**). Notably, the OXPHOS subnetwork can be displayed using different layout options provided by Cytoscape. In our example, we clustered the OXPHOS subnetwork based on different protein complexes (*i.e.*, complex I-V) using Cytoscape *Group Attributes Layout* and *Grid Layout* functions (Error! Reference source not found.**c**).

**Figure 3.**
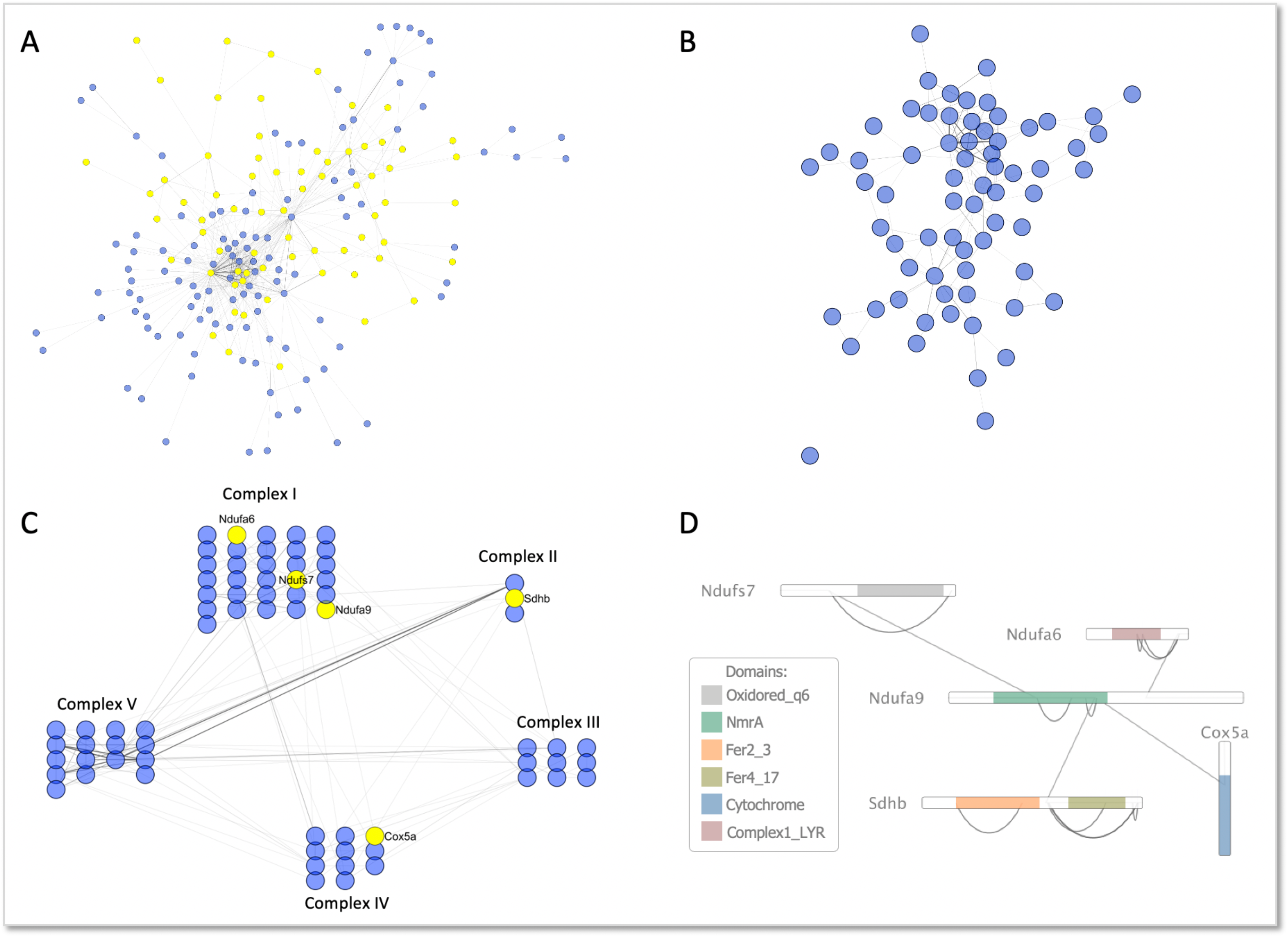
Illustration of XlinkCyNET using a XL-MS-based mitochondrial interactome dataset. (A) The main network cluster of the original dataset displayed using Cytoscape *Preferred Layout*. Highlighted proteins are components of OXPHOS complex. (B) The OXPHOS subnetwork extracted from (A), displayed using Cytoscape *Preferred Layout*. (C) The OXPHOS subnetwork extracted from (A), displayed using Cytoscape *Group Attributes Layout* and *Grid Layout* functions. Highlighted proteins are *Ndufa9* and its first neighboring proteins. (D) The *Ndufa9*-centered subnetwork extracted from (C), displayed in XlinkCyNET layout. Annotated protein domains using Pfam database are shown in colors.

Furthermore, we selected a protein of interest (*Ndufa9* in our example) and its first neighboring proteins using Cytoscape *First Neighbors of Selected Nodes* function and expanded them using XlinkCyNET *apply/restore layout* function (Error! Reference source not found.**d)**. In XlinkCyNET view, residue-to-residue connections and domain annotations are displayed. Cross-link information was loaded in the required XlinkCyNET input file while domain annotations were retrieved using XlinkCyNET *Search for domains* **→** *Pfam* function. For visual simplicity, the XlinkCyNET layout is applied to an extracted subnetwork in our showcase, but it can also be done at any stage of the analysis from the original network.

### XlinkCyNET for elucidating protein complexes topologies: analysis of the XL-MS of the OXPHOS complex I

Next, we illustrate how XlinkCyNET facilitates the analysis of individual protein complex from large scale interactome studie. We focused on mitochondrial OXPHOS complex I, which spans the inner mitochondrial membrane and reaches into the matrix as well as the intermembrane space (IMS). First, we extracted all related nodes and edges of proteins involved in OXPHOS complex I by selecting all proteins labeled as “complex I” in *Node Table* and generated a subnetwork using *New Network From Selected Nodes, All Edges* function (**Figure 4a)**. Next, we applied XlinkCyNET rectangle bar layout to the OXPHOS complex I subnetwork. For domain annotation, we uploaded a manually curated annotation file showing the topologies of proteins based on the high-resolution structure of complex I (PDB: 5LNK). The XlinkCyNET layout clearly shows that cross-links within complex I are confined to specific mitochondrial sub-compartments. That is, residues localized in the IMS and matrix only cross-link to residues in the same sub-compartment (Error! Reference source not found.**b**). This example shows that XlinkCyNET can translate the Cytoscape protein network view into a residue- and domain-specific sub-network, enabling the interrogation of spatial arrangements and interaction sites of protein complexes. These features allow XlinkCyNET to bridge the gap between network visualizations in Cytoscape and molecular visualization systems in structural biology, such as PyMOL (Error! Reference source not found.**a, b**).

**Figure 4.**
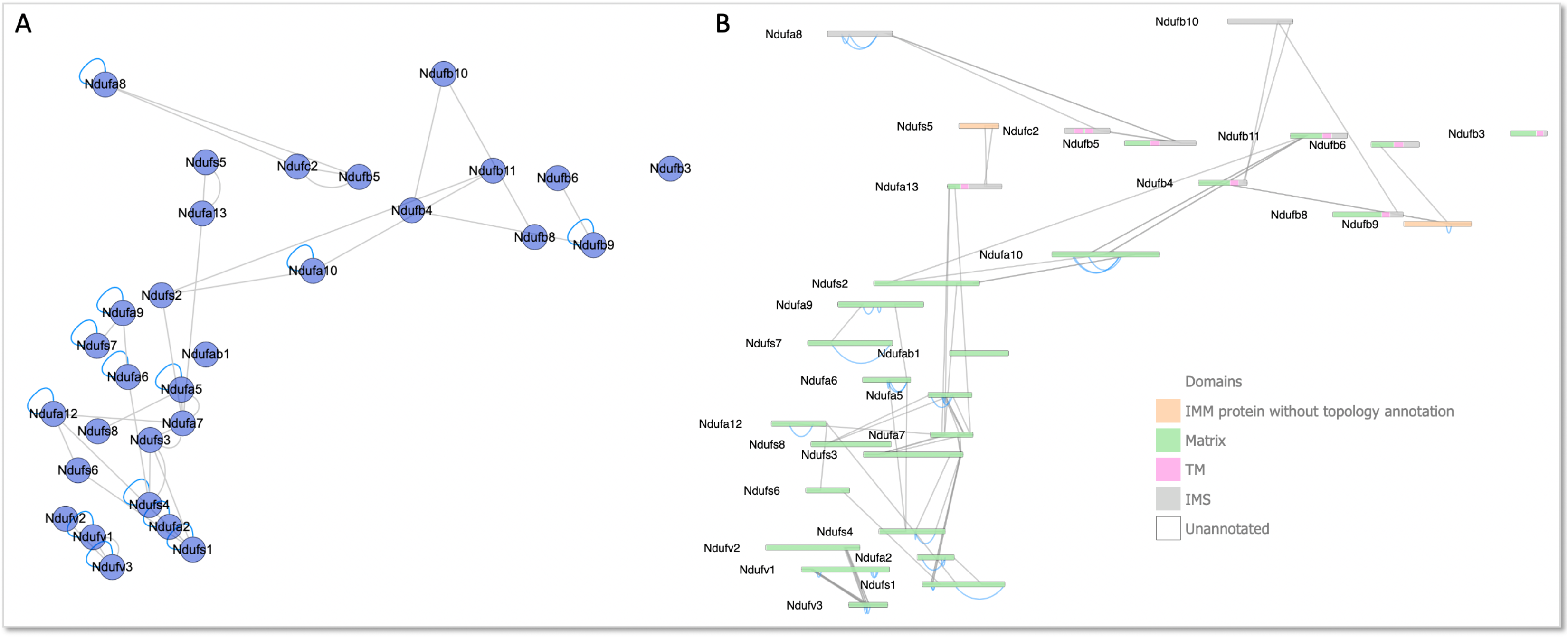
The OXPHOS complex I subnetwork displayed in protein interaction layout (A) and XlinkCyNET layout (B). Domain sub-compartment localizations are indicated in different colors. IMM: inner membrane; TM: transmembrane; IMS: inner membrane space. Proteins are manually arranged based on their positions in the high-resolution structure (PDB: 5LNK).

### XlinkCyNET for revealing interactome changes: analysis of the XL-MS of OXPHOS in native and disrupted mitochondria

Here, we show the usage of XlinkCyNET in profiling interactome changes of mitochondria in two conditions, namely, native and disrupted mitochondria. These two datasets were retrieved from Supplementary Table 1 of the original publication^17^ and formatted to XlinkCyNET-supported *CSV* files using an *in-house* R-script. We first arranged the interaction network of native mitochondria based on protein complexes and functional clusters using Cytoscape *Group Attributes Layout* and *Grid Layout* functions (**Figure 5a**). Next, using another third-party Cytoscape App, namely Copycat Layout^18^ (available at Cytoscape app store), we copied the same protein arrangement from native mitochondria directly to the protein interaction network of disrupted mitochondria (**Figure 5a**). This side-by-side comparison provided an immediate view of the interactome changes of the two conditions. For instance, the interactions between CV (OXPHOS complex V) to mitochondrial matrix proteins (*e.g.*, proteins involved in amino acid metabolism, TCA cycle and fatty acid metabolism) were reduced dramatically. As described in the original paper, the complex I subunit *Ndufa4* underwent substantial rearrangement upon disruption treatment. We therefore selected *Ndufa4* and its first neighboring proteins using *First Neighbors of Selected Nodes* function and applied XlinkCyNET rectangle bar layout to the *Ndufa4*-centered subnetwork (**Figure 5b**). In Cytoscape default layout, we noticed *Ndufa4* obtained a number of additional interaction partners upon disruption treatment (**Figure 5b**). However, the reason for this observation only became obvious when XlinkCyNET layout was applied. Based on the domain annotation illustrating protein sub-compartment localization and topology, we clearly observed that upon disruption treatment, *Ndufa4* mid-localized into mitochondrial matrix side and this resulted in its cross-linking to several matrix-localized proteins (**Figure 5c**).

**Figure 5.**
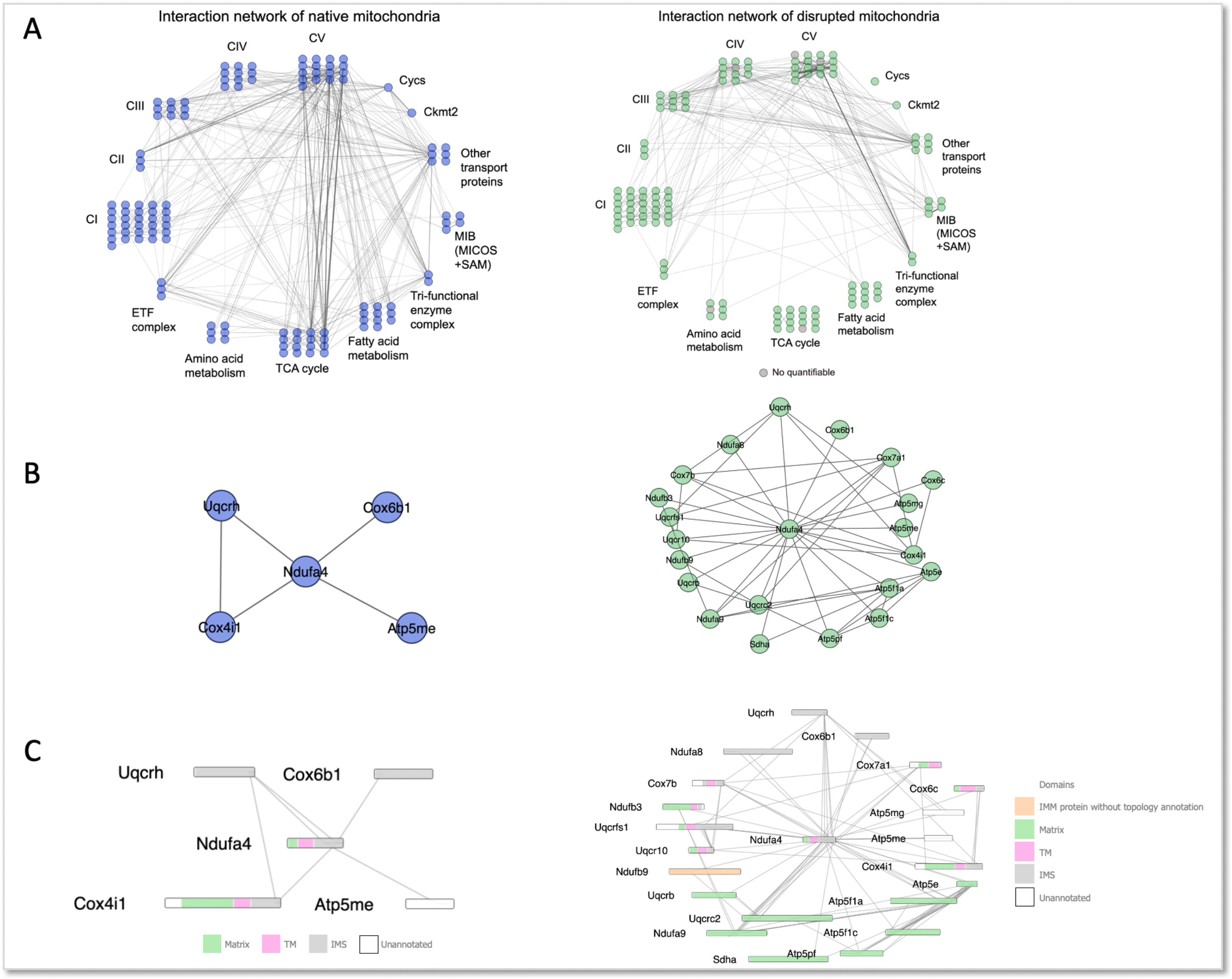
(A) Protein interaction network of native and disrupted mitochondria (showed in blue and green, respectively). (B-C) *Ndufa4*-centered subnetworks of native (left) and disrupted mitochondria (right), shown in Cytoscape default layout (B) and XlinkCyNET layout (C). Domain annotations are colored in the same way as in Figure 4.

## CONCLUSION

We present XlinkCyNET, a fully integrated Cytoscape Java plugin guiding the visualization of XL-MS data in Cytoscape. XlinkCyNET offers optimal presentation of residue-to-residue connections in a rectangular bar style but also supports automatic and manual domain annotation of proteins in the network. Importantly, as a Cytoscape plugin, XlinkCyNET is able to make use of the existing core features embedded in Cytoscape as well as many additional tools provided by third-party plugins available from the Cytoscape app store. In the presented showcases, we illustrated the usage of XlinkCyNET in combination with core features of Cytoscape, such as displaying networks in different layouts and extracting protein- or complex-centered subnetworks to adapt to specific purposes. We also showed examples of using other third-party developed Cytoscape apps, such as Copycat Layout^18^ together with XlinkCyNET for comparison of different networks at sub-domain level. Although not exemplified in our study, importantly, many other Cytoscape apps, such as GeneMANIA^19^, DAVID^20^, BiNGO^21^ and ClueGO^22^ are also compatible with XlinkCyNET. These possibilities offer a wide range of analytical options to XL-MS data, such as functional annotation of proteins and enrichment analysis for set of terms or pathways, thereby greatly expanding the applicability of XlinkCyNET in network analysis. Furthermore, owing to the high-performance of Cytoscape in handling large-scale interaction networks, XlinkCyNET is well suited for network visualization and analysis based on large-scale XL-MS studies.

## ASSOCIATED CONTENT

### Supporting Information

The Supporting Information is available free of charge on the ACS Publications website.

**Table S1**. Examples of protein domain annotation file supported by XlinkCyNET. **Table S2**. The re-formatted *CSV* file for XlinkCyNET input from Supplementary Figure 1 of the Liu *et al.* Paper^17^.

## AUTHOR INFORMATION

### Author Contributions

F.L. designed and supervised the project. D.B.L. developed the software. F.L. and Y.Z. performed early testing of the software and gave feedback. Y.Z. assisted with the XL-MS data analysis. F.L. and D.B.L. wrote the paper. All authors contributed to critical reading of the manuscript.

### Notes

The authors declare no competing financial interest.

1. The web service at http://supfam.org/, for example: http://supfam.org/SUPERFAMILY/cgi-bin/das/up/features?segment=P31946
2. The web service at https://pfam.xfam.org/, for example: https://pfam.xfam.org/protein?entry=P31946&output=xml

## CODE AND DATA AVAILABILITY

XlinkCyNET is an open-source project available at https://github.com/diogobor/XlinkCyNET.

A demonstration video and example data are available at https://www.theliutab.com/software/xlinkcynet.

A detailed protocol is available at protocol exchange^23^ (http://dx.doi.org/10.21203/rs.3.pex-1172/v1).

## ACKNOWLEDGMENT

The authors would like to thank the Leibniz Association with the Leibniz-SAW program (SAW, P70/2018) and the Deutsche Forschungsgemeinschaft (DFG; German Research Foundation, SFB958/Z03) for the financial support for this work.

## REFERENCES

(1) Ma’ayan, A.; Rouillard, A. D.; Clark, N. R.; Wang, Z.; Duan, Q.; Kou, Y. Lean Big Data Integration in Systems Biology and Systems Pharmacology. Trends Pharmacol. Sci. 2014. https://doi.org/10.1016/j.tips.2014.07.001.

(2) Oughtred, R.; Stark, C.; Breitkreutz, B.-J.; Rust, J.; Boucher, L.; Chang, C.; Kolas, N.; O’Donnell, L.; Leung, G.; McAdam, R.; Zhang, F.; Dolma, S.; Willems, A.; Coulombe-Huntington, J.; Chatr-aryamontri, A.; Dolinski, K.; Tyers, M. The BioGRID Interaction Database: 2019 Update. Nucleic Acids Res 2019, 47 (D1), D529–D541. https://doi.org/10.1093/nar/gky1079.

(3) Shannon, P.; Markiel, A.; Ozier, O.; Baliga, N. S.; Wang, J. T.; Ramage, D.; Amin, N.; Schwikowski, B.; Ideker, T. Cytoscape: A Software Environment for Integrated Models of Biomolecular Interaction Networks. Genome Res. 2003, 13 (11), 2498–2504. https://doi.org/10.1101/gr.1239303.

(4) Barsky, A.; Gardy, J. L.; Hancock, R. E. W.; Munzner, T. Cerebral: A Cytoscape Plugin for Layout of and Interaction with Biological Networks Using Subcellular Localization Annotation. Bioinformatics 2007, 23 (8), 1040–1042. https://doi.org/10.1093/bioinformatics/btm057.

(5) Doncheva, N. T.; Morris, J. H.; Gorodkin, J.; Jensen, L. J. Cytoscape StringApp: Network Analysis and Visualization of Proteomics Data. J. Proteome Res. 2019, 18 (2), 623–632. https://doi.org/10.1021/acs.jproteome.8b00702.

(6) Franceschini, A.; Szklarczyk, D.; Frankild, S.; Kuhn, M.; Simonovic, M.; Roth, A.; Lin, J.; Minguez, P.; Bork, P.; von Mering, C.; Jensen, L. J. STRING v9.1: Protein-Protein Interaction Networks, with Increased Coverage and Integration. Nucleic Acids Res 2013, 41 (D1), D808–D815. https://doi.org/10.1093/nar/gks1094.

(7) Leitner, A.; Faini, M.; Stengel, F.; Aebersold, R. Crosslinking and Mass Spectrometry: An Integrated Technology to Understand the Structure and Function of Molecular Machines. Trends in Biochemical Sciences 2016, 41 (1), 20–32. https://doi.org/10.1016/j.tibs.2015.10.008.

(8) Grimm, M.; Zimniak, T.; Kahraman, A.; Herzog, F. XVis: A Web Server for the Schematic Visualization and Interpretation of Crosslink-Derived Spatial Restraints. Nucleic Acids Res 2015, 43 (W1), W362–W369. https://doi.org/10.1093/nar/gkv463.

(9) Combe, C. W.; Fischer, L.; Rappsilber, J. XiNET: Cross-Link Network Maps With Residue Resolution. Molecular & Cellular Proteomics 2015, 14 (4), 1137–1147. https://doi.org/10.1074/mcp.O114.042259.

(10) Lima, D. B.; de Lima, T. B.; Balbuena, T. S.; Neves-Ferreira, A. G. C.; Barbosa, V. C.; Gozzo, F. C.; Carvalho, P. C. SIM-XL: A Powerful and User-Friendly Tool for Peptide Cross-Linking Analysis. Journal of Proteomics 2015. https://doi.org/10.1016/j.jprot.2015.01.013.

(11) Lima, D. B.; Melchior, J. T.; Morris, J.; Barbosa, V. C.; Chamot-Rooke, J.; Fioramonte, M.; Souza, T. A. C. B.; Fischer, J. S. G.; Gozzo, F. C.; Carvalho, P. C.; Davidson, W. S. Characterization of Homodimer Interfaces with Cross-Linking Mass Spectrometry and Isotopically Labeled Proteins. Nature Protocols 2018, 13 (3), 431–458. https://doi.org/10.1038/nprot.2017.113.

(12) de Graaf, S. C.; Klykov, O.; van den Toorn, H.; Scheltema, R. A. Cross-ID: Analysis and Visualization of Complex XL–MS-Driven Protein Interaction Networks. J. Proteome Res. 2019, 18 (2), 642–651. https://doi.org/10.1021/acs.jproteome.8b00725.

(13) UniProt Consortium, T. UniProt: The Universal Protein Knowledgebase. Nucleic Acids Res 2018, 46 (5), 2699–2699. https://doi.org/10.1093/nar/gky092.

(14) Pandurangan, A. P.; Stahlhacke, J.; Oates, M. E.; Smithers, B.; Gough, J. The SUPERFAMILY 2.0 Database: A Significant Proteome Update and a New Webserver. Nucleic Acids Res 2019, 47 (D1), D490–D494. https://doi.org/10.1093/nar/gky1130.

(15) Gough, J.; Karplus, K.; Hughey, R.; Chothia, C. Assignment of Homology to Genome Sequences Using a Library of Hidden Markov Models That Represent All Proteins of Known Structure11Edited by G. Von Heijne. Journal of Molecular Biology 2001, 313 (4), 903–919. https://doi.org/10.1006/jmbi.2001.5080.

(16) Punta, M.; Coggill, P. C.; Eberhardt, R. Y.; Mistry, J.; Tate, J.; Boursnell, C.; Pang, N.; Forslund, K.; Ceric, G.; Clements, J.; Heger, A.; Holm, L.; Sonnhammer, E. L. L.; Eddy, S. R.; Bateman, A.; Finn, R. D. The Pfam Protein Families Database. Nucleic Acids Res. 2012, 40 (Database issue), D290–301. https://doi.org/10.1093/nar/gkr1065.

(17) Liu, F.; Lössl, P.; Rabbitts, B. M.; Balaban, R. S.; Heck, A. J. R. The Interactome of Intact Mitochondria by Cross-Linking Mass Spectrometry Provides Evidence for Coexisting Respiratory Supercomplexes. Molecular & Cellular Proteomics 2018, 17 (2), 216–232. https://doi.org/10.1074/mcp.RA117.000470.

(18) Settle, B.; Otasek, D.; Morris, J. H.; Demchak, B. Copycat Layout: Network Layout Alignment via Cytoscape Automation. F1000Res 2018, 7, 822. https://doi.org/10.12688/f1000research.15144.1.

(19) Montojo, J.; Zuberi, K.; Rodriguez, H.; Bader, G. D.; Morris, Q. GeneMANIA: Fast Gene Network Construction and Function Prediction for Cytoscape. F1000Res 2014, 3, 153. https://doi.org/10.12688/f1000research.4572.1.

(20) Dennis, G.; Sherman, B. T.; Hosack, D. A.; Yang, J.; Gao, W.; Lane, H. C.; Lempicki, R. A. DAVID: Database for Annotation, Visualization, and Integrated Discovery. Genome Biology 2003, 4 (5), P3. https://doi.org/10.1186/gb-2003-4-5-p3.

(21) Maere, S.; Heymans, K.; Kuiper, M. BiNGO: A Cytoscape Plugin to Assess Overrepresentation of Gene Ontology Categories in Biological Networks. Bioinformatics 2005, 21 (16), 3448–3449. https://doi.org/10.1093/bioinformatics/bti551.

(22) Bindea, G.; Mlecnik, B.; Hackl, H.; Charoentong, P.; Tosolini, M.; Kirilovsky, A.; Fridman, W.-H.; Pagès, F.; Trajanoski, Z.; Galon, J. ClueGO: A Cytoscape Plug-in to Decipher Functionally Grouped Gene Ontology and Pathway Annotation Networks. Bioinformatics 2009, 25 (8), 1091–1093. https://doi.org/10.1093/bioinformatics/btp101.

(23) Borges Lima, D.; Zhu, Y.; Liu, F. Using XlinkCyNET to Visualize Protein Cross-Links in Cytoscape Software. Protocol Exchange. https://doi.org/10.21203/rs.3.pex-1172/v1.

